# The capsule of *Cryptococcus neoformans* modulates phagosomal pH through its acid-base properties

**DOI:** 10.1101/390328

**Authors:** Carlos M. De Leon-Rodriguez, Man Shun Fu, M. Osman Corbali, Radames J.B. Cordero, Arturo Casadevall

## Abstract

Phagosomal acidification is a critical cellular mechanism for the inhibition and killing of ingested microbes by phagocytic cells. The acidic environment activates microbicidal proteins and creates an unfavorable environment for the growth of many microbes. Consequently, numerous pathogenic microbes have developed strategies for countering phagosomal acidification through various mechanisms that include interference with phagosome maturation. The human pathogenic fungus *Cryptococcus neoformans* resides in acidic phagosome after macrophage ingestion that actually provides a favorable environment for replication since the fungus replicates faster at acidic pH. We hypothesized that the glucuronic acid residues in the capsular polysaccharide had the capacity to affect phagosome acidity through their acid-base properties. A ratiometric fluorescence comparison of imaged phagosomes containing *C. neoformans* to those containing beads showed that the latter were significantly more acidic. Similarly, phagosomes containing non-encapsulated *C. neoformans* cells were more acidic than those containing encapsulated cells. Acid-base titrations of isolated *C. neoformans* polysaccharide revealed that it behaves as a weak acid with maximal buffering capacity around pH 4-5. We interpret these results as indicating that the glucuronic acid residues in the *C. neoformans* capsular polysaccharide can buffer phagosomal acidification. Interference with phagosomal acidification represents a new function for the cryptococcal capsule in virulence and suggests the importance of considering the acid-base properties of microbial capsules in the host-microbe interaction for other microbes with charged residues in their capsules.

**Importance:** *Cryptococcus neoformans* is the causative agent of cryptococcosis, a devastating fungal disease that affects thousands of individuals worldwide. This fungus has the capacity to survive inside phagocytic cells, which contributes to persistence of infection and dissemination. One of the major mechanisms of host phagocytes is to acidify the phagosomal compartment after ingestion of microbes. This study shows that the capsule of *C. neoformans* can interfere with full phagosomal acidification by serving as a buffer.

## Introduction

Ingestion of microbes by phagocytic cells results in the formation of a new organelle called the phagolysosome, a membrane-bound compartment where microorganisms are subjected to a variety of antimicrobial compounds such as oxidative radicals and microbicidal proteins. The process of phagolysosome formation results from a complex cellular choreography that includes phagosome acidification, resulting in microbial inhibition by creating an unfavorable environment and the activation of various microbicidal compounds. Consequently, diverse pathogenic microbes have developed mechanisms to interfere with phagosomal acidification. For example, the bacterium *Mycobacterium tuberculosis* (1), the fungus *Histoplasma capsulatum* (2), and the parasite *Leishmania donovani* (3) each avoids phagosomal acidification by interfering with the process of phagosome maturation, which reduces the presence of the vesicular proton-ATPase from phagosome. Hence, modulation of phagosomal acidification by microbes ingested by phagocytic cells and the mechanisms for such effects are topics of great interested and research activity in the field of microbial pathogenesis research.

*Cryptococcus neoformans* is a facultative intracellular pathogenic yeast (4) that is a major cause of meningoencephalitis in individuals with impaired immunity (5). In contrast to many other facultative intracellular pathogens, this fungus resides in an acidic phagosome after ingestion by macrophages (6). Despite residence in acidic phagosome, there is evidence that the *C. neoformans* modulates some aspects of phagosomal maturation, including full phagosomal acidification, although the mechanisms for this effect have not been fully elucidated (7, 8). In fact, for *C. neoformans,* acidification has been viewed as favoring intracellular growth since this fungus replicates faster in acidic environments (9). Its survival inside the phagosome is believed to result from its ability to withstand oxidative bursts (10), damage the phagosomal membranes (11) and damage critical host cell homeostasis (12) rather than interference with phagosomal maturation, although the relative contributions to the overall outcome of intracellular survival remain to be determined.

Recently, we reported a new role for *C. neoformans* urease in modulating phagosomal pH (13). Urease positive *C. neoformans* strains hydrolyzed urea to ammonia resulting in pleiotropic changes to the cryptococcal macrophage interaction that included higher phagosomal pH, delayed intracellular growth, and enhanced non-lytic exocytosis (13). *C. neoformans* is unusual among intracellular pathogens in that it grows faster at lower pH resulting in faster replication inside phagolysosomes than in the extracellular medium (9). Loss of phagosomal integrity is associated with reduced acidity in that compartment and the triggering of macrophage death (14). Hence, the extent of phagosomal acidification is an important variable, which can favor the microbe or the host cell depending on the state of the interaction (13, 14).

One of the most striking characteristics of *C. neoformans* as a pathogenic microbe is that it is surrounded by a large polysaccharide capsule that is a critical determinant of virulence (15). The capsule functions in virulence by interfering with phagocytosis and immune responses (15, 16). The capsule is also thought to play a major role in intracellular survival by quenching free radical fluxes in the phagosome (10). The major capsular polysaccharide is glucuronoxylomannan (GXM), which is composed of a mannose backbone with xylose and glucuronic acid substitutions (17). The presence of glucuronic acid residues in cryptococcal polysaccharide imparts a negative charge to the capsule (18), that is believed to contribute to protection against phagocytosis. In addition, those glucuronic acid residues can be anticipated to impart considerable acid-base properties to the cryptococcal GXM. In our recent study on the role of phagosomal membrane integrity, we observed that even though apoptotic cells had higher phagolysosomal pH, loss of membrane integrity was not associated with complete loss of acidity, which we hypothesized was due to the acid-based properties of the capsule (14). In contrast, for *Candida albicans,* which lacks a polysaccharide capsule and hence has no comparable buffering capacity, phagosome permeabilization resulted in luminal alkalinization (19). In this study, we formally tested that hypothesis and present evidence that the capsule of *C. neoformans* interferes with full phagosome acidification. These findings establish a new mechanism for microbial modulation of phagosomal pH and imply a new role for the capsule in cryptococcal virulence.

## Materials and Methods

### Yeast culture

*C. neoformans* serotype A strain H99 and the acapsular mutant *cap59* were used for all experiments. Cells were grown from frozen stocks in Sabouroaud dextrose liquid media at 30°C under agitation (180 rpm) for 2 d.

### Coating of acapsular mutant with GXM

For the formation of the proto-capsule we follow previously published methods (20). Briefly, the supernatant of overnight culture of H99 was cleared by centrifugation and filtered using a 0.8 μm syringe filter. An overnight culture of 1 x 10^7^ cells/mL *cap59* acapsular strain was then incubated with 100, 10 or 1 μL of H99 cleared supernatant (conditioned media) in a total 1 mL medium with rotation for 1 h at room temperature. Images were acquired using the Olympus AX70 microscopy (Olympus, Center Valley, PA) with objective 40x to visualize the formation of the proto-capsule, which was labeled by Oregon green 488 conjugated 18B7 monoclonal antibody.

### Measurement of phagosomal pH

Phagolysosomal pH was measured using ratiometric fluorescence imaging involving the use of pH-sensitive probe Oregon green 488. Oregon green 488 was first conjugated to monoclonal antibody 18B7 using Oregon Green 488 Protein Labeling Kit (Molecular Probes, Eugene, OR), as described (13). The Oregon Green 488 dye has a succinimidyl ester moiety that reacts with primary amines of proteins to form stable dye-protein conjugates. The labeling procedure is according to the manufacture’s instruction. BMDM were plated (4 × 10^5^ cells/well) on 24-well plate with 12 mm circular coverslip coated with 100 μg/mL poly-D-lysine. Cells were cultured with completed BMEM medium containing 0.5 μg/mL LPS and 100 U/mL IFN-γ and then incubated at 37 °C with 9.5 % CO_2_ overnight. For infection, H99 and *cap59* strains (8 × 10^6^ cells/mL) were incubated with 10 μg/mL Oregon green conjugated mAb18B7 for 15 min. Macrophages were then infected with Oregon green conjugated 18B7-opsonized yeast in 4 × 10^5^ cells per well. Cells were centrifuged immediately at 270 *g* for 1 min and culture were incubated at 37°C for 10 min to allow phagocytosis. Extracellular cryptococcal cells or beads were removed by washing three times with fresh medium. Samples on coverslip were collected at 24 h after phagocytosis by washing twice with pre-warmed HBSS. Annexin V Alexa Fluor 555 staining was performed as manufacturer’s instructions (Invitrogen, Carlsbad, CA). The coverslip was then placed upside down on MatTek petri dish (35 mm; 10 mm diameter microwell; MatTek, Ashland, MA) with the Annexin V binding buffer in the microwell. Images were taken by using Olympus AX70 microscopy (Olympus, Center Valley, PA) with objective 40x at dual excitation 440 nm and 488 nm for Oregon green, 550 mm for Annexin V and bright field. Images were acquired and analyzed using MetaFluor Fluorescence Ratio Imaging Software (Molecular Devices, Downingtown, PA). Relative phagolysosomal pH was determined based on the ratio of 488 nm/440 nm. The relative pH was converted to absolute pH by obtaining the standard curve in which the images are taken as above but intracellular pH of macrophage was equilibrated by adding 10 μM nigericin in pH buffer (140 mM KCl, 1 mM MgCl_2_, 1 mM CaCl_2_, 5 mM glucose, and appropriate buffer ≤ pH 5.0: acetate-acetic acid; pH 5.5-6.5: MES; ≥ pH 7.0: HEPES. Desired pH values were adjusted H using either 1M KOH or 1M HCl). Buffers were used at pH 3-7.5 using 0.5-pH unit increments.

### Biotinylation of cells

Approximately 1 × 10^6^ cryptococcal cells were biotinylated using EZ Link-Sulfo-NHS-biotin (21217, ThermoScientific, Rockford, IL). Overnight cultures were washed three times with PBS pH 8.0, and diluted in PBS pH 8.0 to 1 × 10^6^ cells/ml. EZ Link-Sulfo-NHS-biotin in 2 mM was added in the cell samples, and incubated for 30 min at room temperature. Cells were then washed three times with PBS with 100 mM glycine to remove excess biotin reagent and byproduct. After biotinylation, cells were labeled with 5 μg/ml Oregon Green conjugate of NeutrAvidin biotic-binding protein (A6374, ThermoFisher Scientific) with rotation for 1 h at room temperature.

### GXM isolation

Soluble GXM was obtained from culture supernatants of encapsulated cells by ultrafiltration (21, 22). Briefly, culture supernatants were collected by centrifugation (6000 x *g,* 15 min, 4°C) and filtered using 0.22 μm vacuum driven disposable bottle-top filter (MilliPore) to ensure clearing of cells and other large debris. The cleared supernatant was ultra-filtered sequentially in an Amicon ultrafiltration cell (Millipore, Danvers, MA) using membranes of different nominal molecular weight limits (100 and 10 kDa). After filtrating using a 100 KDa, the flow-through was again filtered through 10 kDa. On each filtration step, GXM can be recovered from the surface of membranes in the form of a viscous gel. This process yields GXM fractions of >100 kDa and 100-10 kDa that were then dialyzed against ultrapure water, lyophilized and store until use.

### Acid-based titrations

Forty milliliter solutions of 3 and 10 mM sodium D-glucuronate (Sigma G8645) were titrated against 0.1 M HCl in 40 mL glass beakers. Since the pH of ultrapure water was acidic ~pH 5, it was adjusted to pH 7 using NaOH before preparing the glucoronate solutions. GXM solutions were prepared by dissolving the lyophilized GXM fraction 10-100 kDa in ultrapure water at 1 mg/mL. Since GXM molecules exhibit wide size distributions, from 1 to >100 kDa, we used the 10-100 kDa fraction to narrow the range of molecular mass. Total monosaccharide concentration in GXM solutions were determined by the Dubois method (AKA phenol-sulfuric acid assay (23). After phenol sulfuric assay, GXM solutions were diluted to 2.4 mM total monosaccharide concentration and 3 mL volumes were titrated against 0.01 M HCl in 10 mL glass beakers. Titrations were conducted in beakers placed inside a water bath equilibrated at 20°C temperature under constant stirring. Changes in pH were recorded using an Accumet pH meter.

### Calculation of theoretical acid-base titration curve

To calculate the theoretical acid-base titration curve for sodium D-glucuronate (NaA) we assumed that positive charges and negative charges are equal in an ideal solution: [*Na*^+^] + [*H*^+^] = [*OH*^−^] + [*A*^−^] + [*Cl*^−^], such that *NaA* ↔ *Na*^+^ + *A*^−^ and *NaA* + *HCl* ↔ *HA* + *NaCl*. Such that: [*Na*+] = [*A*^−^] + [*HA*], since *C_HA_* = [*HA*] + [*A*^−^]. Also, *C_HCl_* = [*Cl*^−^] + [*HCl*] and since HCl totally dissolves, then *C_HCl_* = [*Cl*].

Hence, the equation becomes;

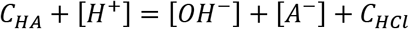

At this point, we can turn this equation into a third-degree polynomial, with [*H*^+^] being the unknown.

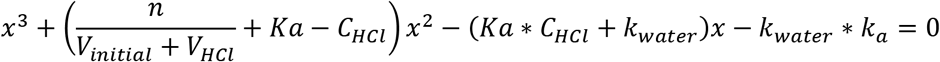

while pKa = 3.28 (Glucuronic Acid), K_water_=0.681*10^−14^ at 20°C and n is initial amount of moles of sodium D-glucuronate we use for our solution. Biggest positive root of this third degree polynomial provided us the theoretical [*H*^+^] value after a certain moles of acid added.

### Statistical analysis

All statistical analyses were performed by using One-way ANOVA, followed by Tukey’s or Dunnett’s multiple-comparison test.

## Results

### pH of phagosomes containing beads and *C. neoformans*

Ingested *C. neoformans* resides in a mature acidic phagosome (6). However, the extent to which *C. neoformans* modulates the pH of the cryptococcal phagosome is unknown. A comparison of the pH of phagosomes containing inert beads with phagosomes containing *C. neoformans* cells showed that the latter were significantly less acidic (Figure 1). On average the pH of phagosomes containing inert beads was 4.22 ± 0.45 (n = 40) at 3 h, which corresponded to a 0.65 pH unit difference (p < 0.0001 by one-way ANOVA and Tukey’s multiple-comparison test). To ascertain whether this higher pH was the result of active pH modulation by *C. neoformans* we compared the pH of phagosomes containing live and dead *C. neoformans* cells. Comparison of the average pH in phagosomes containing live and dead *C. neoformans* cells revealed average values of 4.87 ± 0.58 (n = 62) and 4.78 ± 0.14 (n = 43) at 3 h, respectively, (*P* = 0.638 by one-way ANOVA and Tukey’s multiple-comparison test). Phagosomes containing live and dead cells had comparable pH to those having live cells, suggesting that the pH modulation in the phagosome is not the result of secretion of basic compounds by *C. neoformans* (Figure 1).

**Figure 1.**
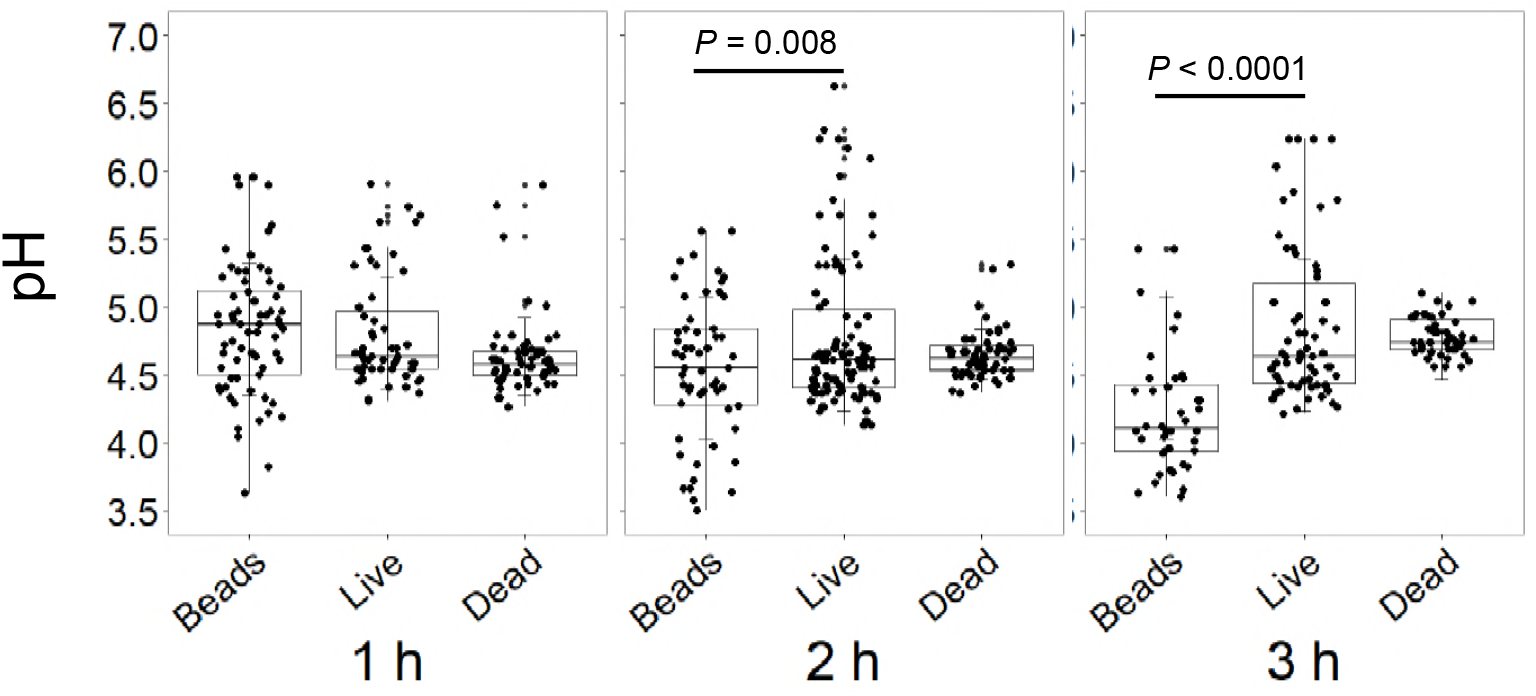
pH of phagosomes containing beads, heat-killed and live *C. neoformans.* Phagolysosomal pH was measured by using Oregon-green dual-excitation ratio fluorescence imaging at indicated time point. Each dot represents pH of individual phagolysosomes. Data in are from one representative experiment. Comparable results were obtained from two additional independent experiment. *P* values by one-way ANOVA with Tukey’s multiple comparison test.

### Phagosomal pH of encapsulated cells is higher than of non-encapsulated cells

To test the hypothesis that pH modulation was the result of the acid-base properties of the *C. neoformans* we sought to compare the phagosomal pH for encapsulated and non-encapsulated cells. However, this presented the practical problem that non-encapsulated cells could not be opsonized through the FcR since they lacked a capsule that would bind GXM-binding antibody. Opsonizing encapsulated cells with antibody and nonencapsulated cells with complement was not considered acceptable since the two opsonins are very different. We tried to label cryptococcal cells with EZ-Link NHS-Biotin and Oregon Green^®^ 488 conjugated NeutrAvidin®, and the phagocytosis was performed in the use of guinea pig complement. However, the signal of Oregon green was not stable in the phagosome and after 24 h it was lost completely, which we attribute to dye degradation from a combination of the low phagosomal pH and the oxidative burst (24). We also have tried to measure the phagolysosomal pH with heat-killed cryptococcal cells labeled with NHS-Biotin but the labeling did not work with heat-killed cells. Hence, we resorted to coating non-encapsulated cells with encapsulated *C. neoformans* conditioned media, which results in the attachment of soluble polysaccharide to the surface of non-encapsulated cells to create a proto-capsule that would allow antibody-mediated opsonization (Figure 2). The phagosomal pH of naturally encapsulated *C. neoformans* cells was significantly larger than that of non-encapsulated cells containing an artificial capsule (Figure 3).

**Figure 2.**
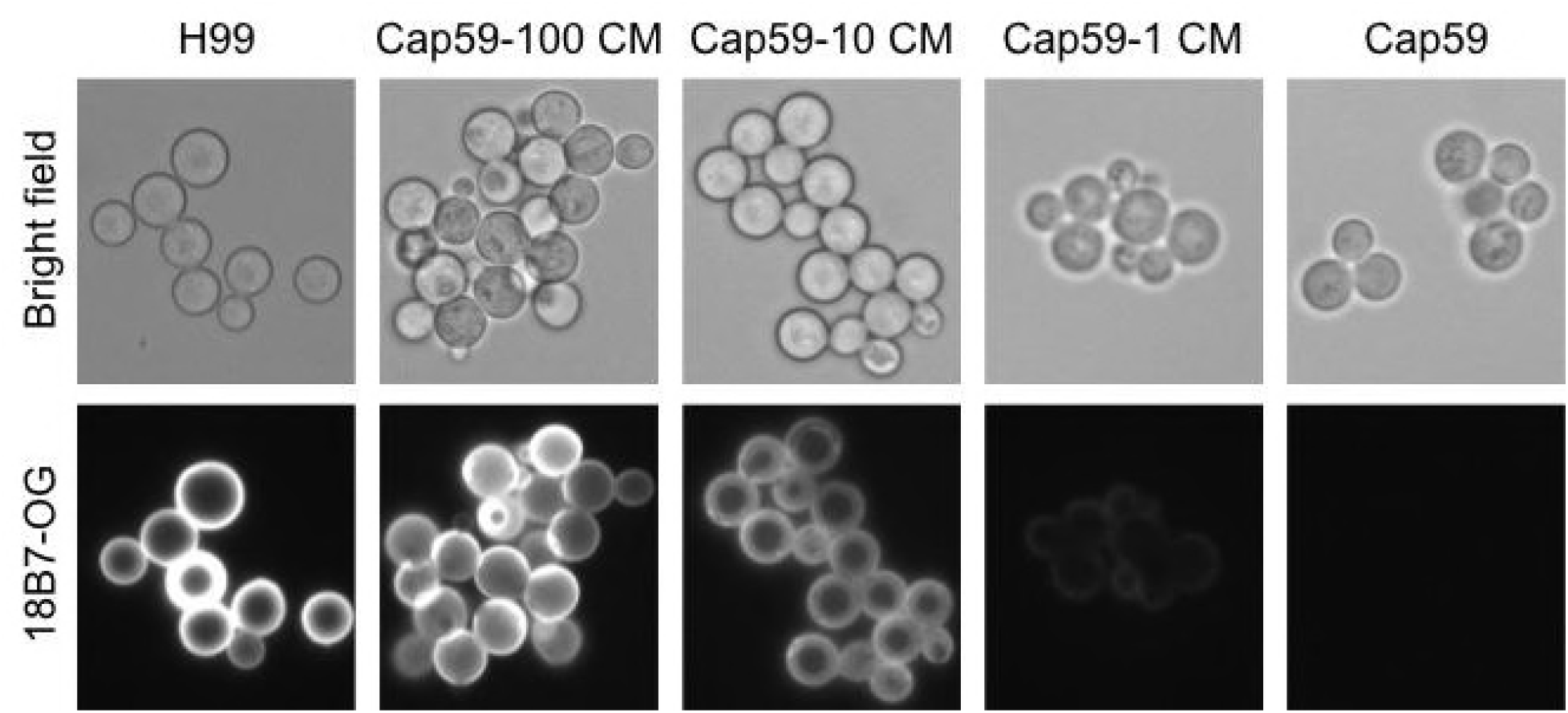
Binding of capsular polysaccharide to the non-encapsulated *C. neoformans* strain *cap59.* Capsule material (CM) release into the media by *C. neoformans* was incubated at different concentrations with *cap59* to form a proto-capsule around these cells. Subsequently, the cells were incubated with the monoclonal anti body 18B7 previously conjugated with Oregon Green (18B7-OG). Bright field (top) and immunofluorescence (bottom) images are shown of *C. neoformans* H99 the acapsular strain *cap59* incubate with 100 ul, 10 ul, 1 ul of CM and *cap59* alone. The magnification for this figure is 40X.

**Figure 3.**
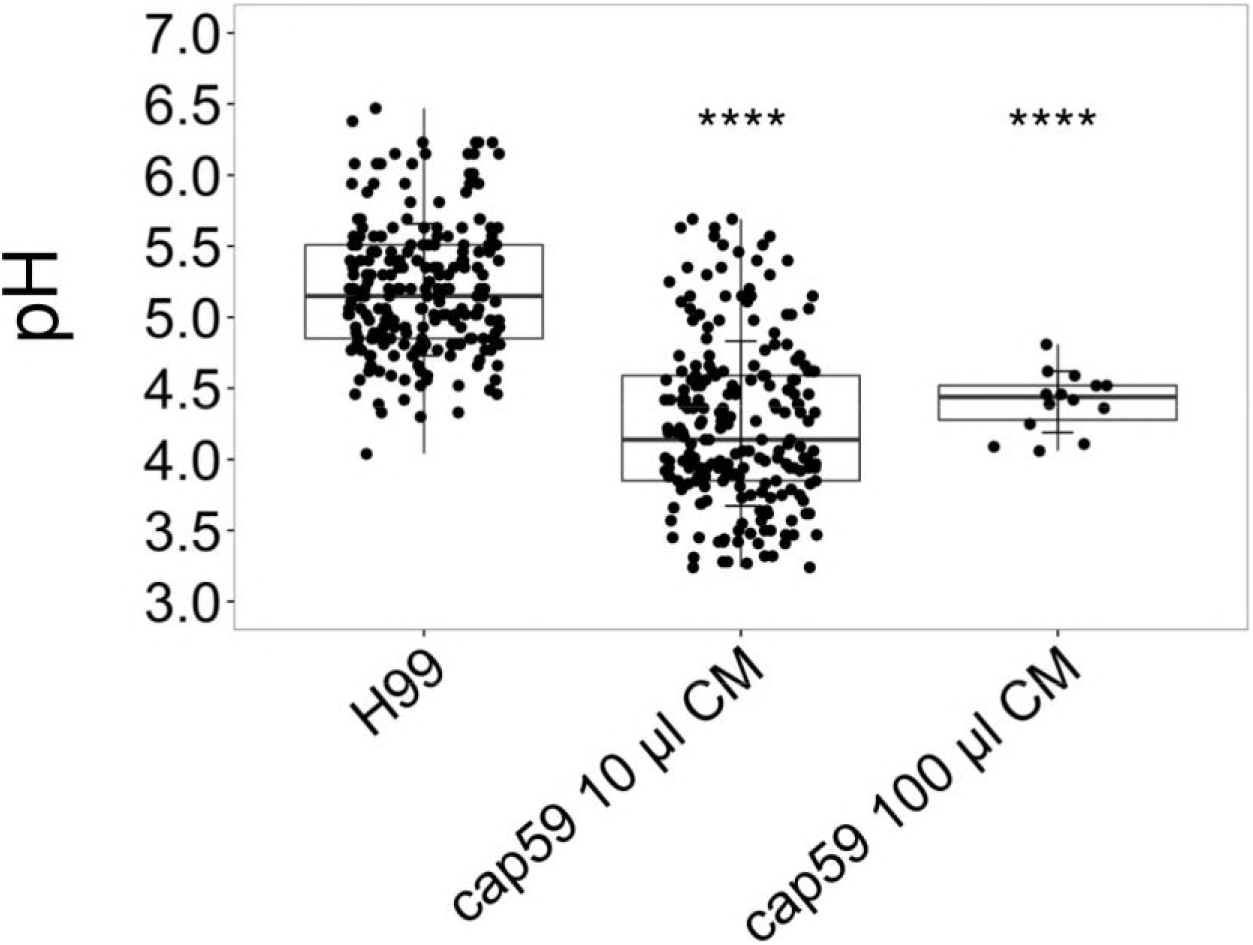
pH measurement of phagolysosomal-containing encapsulated and proto-encapsulated *C. neoformans.* Macrophages were infected with encapsulated (H99) and *cap59* previously incubated with 100 and 10 μL of conditioned media (CM) to form a proto-capsule. Opsonization was antibody-mediated using 18B7 bound to Oregon Green. The pH value was 5.19 in H99- containing-phagolysosomes which was less acidic than phagosomes containing *cap59* incubated with 100 and 10 μL of CM, which had pH values of 4.40 and 4.25 respectively. One-way ANOVA with Dunnett’s multiple comparison test, **** p < 0.0001.

### Acid-base properties of glucuronic acid and GXM

Glucuronic acid is an organic weak acid with a relatively high pKa. We titrated a sodium salt of glucuronic acid (sodium D-glucuronate) with HCl and calculated a pKa in the range of 2.5-3.11 at the beginning of our titrations (corresponding to 0.23-20 μmoles of titrant) (Figure 4A). These values are comparable to the reported pKa values 2.9 (25), 2.8-2.9 from ^13^C-nuclear magnetic spectroscopy (26) and 3.28 measured from standard acid-based titrations (27). Consequently, the presence of glucuronic acid in solution confers considerable buffering capacity such that for a 10 mM solution to change from pH 7 to pH 4, it requires almost 10 times the acid required to achieve the same pH reduction as in a pure water solution. Similarly, the presence of GXM in water provided considerable buffering capacity at around pH 5, which is close to the final pH in cryptococcal phagosomes (Figure 4B). Considering the known polyelectrolyte nature and polydispersity of GXM preparations in terms of molecular mass 16278213, 21208301, together with the mild inflection point, it is problematic to determine a pKa value for GXM.

**Figure 4.**
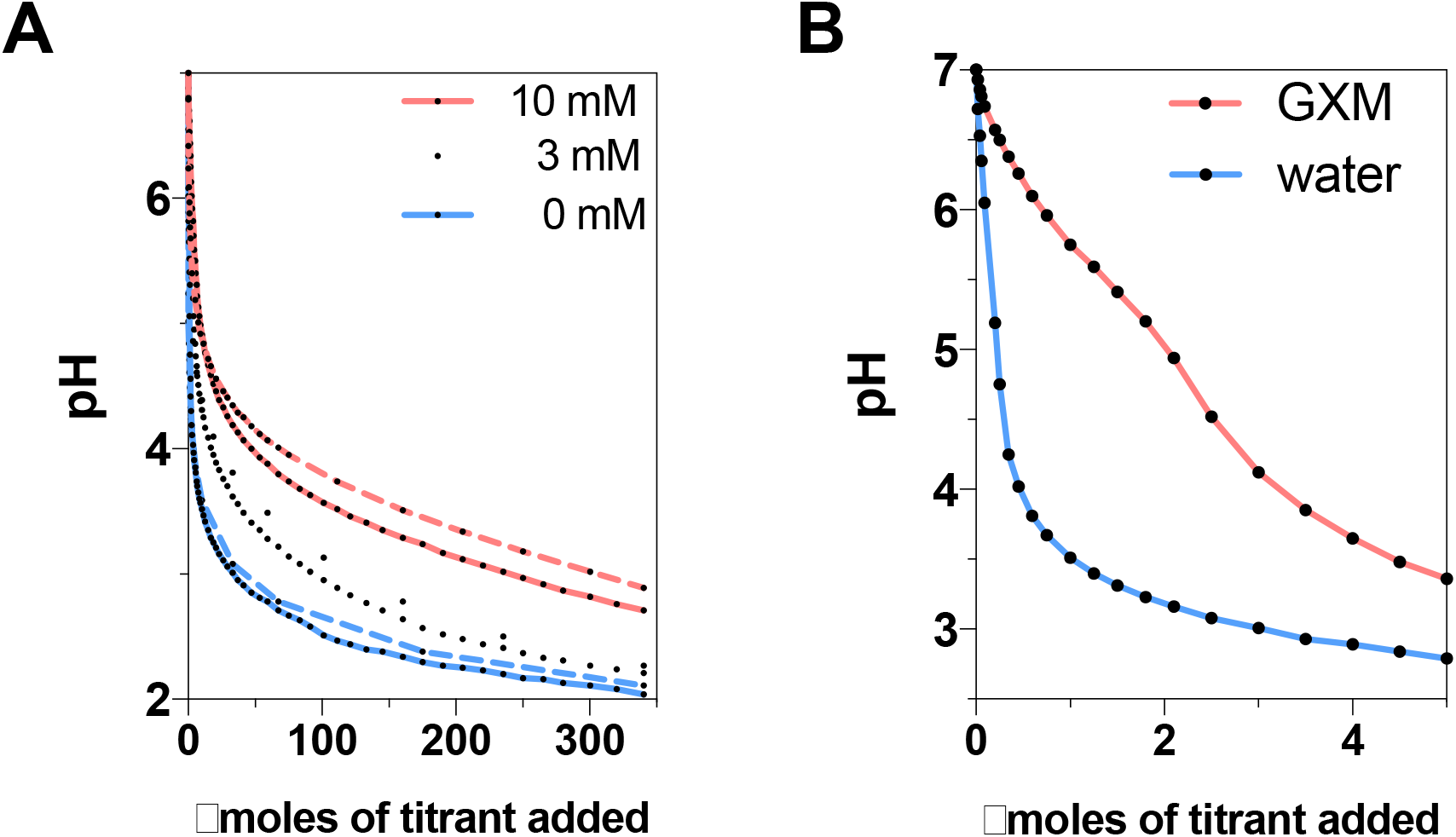
Titration of sodium D-glucuronate and GXM against HCl. (A) Changes in pH of sodium D-glucuronate solutions at 3 and 10 mM as a function of micromoles of titrant added. Initially, sodium D-glucuronate solutions show a rapid pH change by acidic titration. Near to pKa level (pH~3.28) it shows the best buffering capacity compared to pure water. Theoretical curves (dashed lines) for glucuronic acid also reveal a similar tendency of rapid pH change and buffering capacity. (B) Change in pH of GXM solution as a function of micromoles of titrant added. GXM provides a substantial amount of buffering capacity around pH 5 compared to ultrapure water.

## Discussion

Many pathogenic microbes express polysaccharide capsules that are essential for virulence. Microbial capsules are directly antiphagocytic and efficient ingestion of encapsulated microbes by phagocytic cells usually require antibody- or complement-derived opsonins. However, many microbial capsules are composed of polysaccharides that are poorly immunogenic that often fail to induce strong responses. Although microbial capsules are generally thought to contribute to virulence by resisting ingestion and killing by host phagocytic cells, there is evidence that capsules mediate other functions that contribute to pathogenesis and that some of these effects are mediated by capsule ionic charge. For instance, negatively charged capsular polysaccharides of gram negative bacteria can bind cationic microbicidal peptides and protect bacterial cells (28). On the other hand, the positively charged modifications in *Streptococcus pneumonia* capsular polysaccharide can predispose bacterial cells to enhanced killing by alpha-defensins (29). Among *Cryptococcus* spp., a comparison of glucuronic acid residue content among a non-pathogenic and pathogenic spp. revealed that the latter had higher content of this charged residue (30). Hence, while there is considerable evidence that microbial surface charge can play an important role in pathogenesis through a variety of mechanisms their role in phagosomal pH modulation has not been investigated.

It is axiomatic that microbial capsules containing weak acid and basic residues will exhibit acid-base properties that are reflective of the capacity of these residues to function as proton donors and acceptors. In this regard, the cryptococcal capsule is composed of a repeating mannose triad, of which each includes one glucuronic acid residues, that in turn confers upon the polysaccharide and the resulting capsule a negative charge (18). Our results confirm the theoretical deduction that the *C. neoformans* capsular GXM buffers the phagolysosomal pH. While titration of sodium salt of glucuronic acid (Sodium D-Glucuronate) shows a rapid initial decrease in pH during titration, GXM provides a buffer effect around pH 5. This is not surprising since pKa of glucuronic acid is 3.28 and solution of sodium D-glucuronate will exert its best buffering effect near the pKa. However, with readily available acidic and basic residues of glucuronic acid (*A*^−^ and *HA*) covalently bound to the GXM polysaccharide backbone, these residues could experience different electronic milieu that could modify their ionic properties. Furthermore, GXM molecules are large and structurally complex and consequently not all glucuronic acids may be equally exposed to the solvent and in a position to donate or accept hydronium ions equally. Despite these differences, GXM retains considerable weak acid properties that would confer a maximum buffering capacity at the pH range of approximately 4-5, which corresponds to the optimal pH for *C. neoformans* growth (13).

Ingested dead encapsulated *C. neoformans* cells resided in a phagosome that had a higher pH than inert beads. This suggested that *C. neoformans* cells contained anionic groups that could buffer hydronium ions in the phagosome resulting in a higher pH. To ascertain the contribution of the capsule to this effect we compared the phagosomal pH of *C. neoformans* encapsulated cells to acapsular cells coated with a proto-capsule that would allow both to be opsonized with IgG1 through the same Fc receptors.

We recognize that in adding GXM to acapsular cells to create a proto-capsule that permitted antibody-mediated opsonization meant that we undermined any comparison involving GXM acid-base effects since this maneuver introduced some polysaccharide into the phagosome. We reasoned that this handicap was outweighed by the fact that we could not be certain that other methods of opsonization would result in comparable phagosome and the fact that natural capsules are much larger than artificial capsules meant that there would still be much more polysaccharide in encapsulated cryptococcal phagosomes. Despite this handicap, we were able to measure a difference in phagosomal pH between encapsulated and acapsular strains consistent with a strong acid buffering capacity by the polysaccharide capsule.

In summary, the presence of glucuronic acid residues in the *C. neoformans* capsule makes the polysaccharide a weak acid capable of modulating pH in the phagosome. Our experimental observations are consistent with the expected acid-base properties of the capsule based on its sugar residue composition. The fact that the polysaccharide capsule of *C. neoformans* is large brings considerable GXM mass into the phagosome with the potential to mediate considerable buffering capacity. Given that *C. neoformans* has optimal growth rate at acidic pHs (13), the acid-base properties of the capsule can be expected to promote fungal cell survival in the phagosome by its buffering capacity during conditions of both phagosomal acidification and phagosomal membrane leakage. This mechanism for phagosomal pH modulation based on acid-base properties is very different from used by other intracellular pathogens that modulate pH by interfering with phagosome maturation. Our observations suggest that other microbes with charged microbial capsules could also modulate phagosomal acidification through their acid-based electrolyte properties.

## Acknowledgements

AC was is supported in part by 5R01HL059842, 5R01AI033774, 5R37AI033142, and 5R01AI052733

